# PatternMarkers & GWCoGAPS for novel data-driven biomarkers via whole transcriptome NMF

**DOI:** 10.1101/083717

**Authors:** Genevieve L Stein-O’Brien, Jacob L Carey, Wai-shing Lee, Michael Considine, Alexander V Favorov, Emily Flam, Theresa Guo, Sijia Li, Luigi Marchionni, Thomas Sherman, Shawn Sivy, Daria A Gaykalova, Ronald D McKay, Michael F Ochs, Carlo Colantuoni, Elana J Fertig

## Abstract

**Summary:** Non-negative Matrix Factorization (NMF) algorithms associate gene expression with biological processes (e.g., time-course dynamics or disease subtypes). Compared with univariate associations, the relative weights of NMF solutions can obscure biomarkers. Therefore, we developed a novel PatternMarkers statistic to extract genes for biological validation and enhanced visualization of NMF results. Finding novel and unbiased gene markers with PatternMarkers requires whole-genome data. However, NMF algorithms typically do not converge for the tens of thousands of genes in genome-wide profiling. Therefore, we also developed Genome-Wide CoGAPS Analysis in Parallel Sets (GWCoGAPS), the first robust whole genome Bayesian NMF using the sparse, MCMC algorithm, CoGAPS. This software contains analytic and visualization tools including a Shiny web application, patternMatcher, which are generalized for any NMF. Using these tools, we find granular brain-region and cell-type specific signatures with corresponding biomarkers in GTex data, illustrating GWCoGAPS and patternMarkers ascertainment of data-driven biomarkers from whole-genome data.

**Availability:** PatternMarkers & GWCoGAPS are in the CoGAPS Bioconductor package (3.5) under the GPL license.

## 1 Introduction

Numerous high-throughput studies link gene expression changes to biological processes (BPs) including regulatory networks and the cell signaling processes. Previously shown effective at deconvoluting multiplexed regulation and gene reuse in BPs (Trendafilov and Unkel, 2011; Kossenkov and Ochs, 2009; Ochs and Fertig, 2012), NMF algorithms have identified genes associated with yeast cell cycle and metabolism, cancer subtypes, and perturbations to cellular signaling in cancer (Li and Ngom, 2013; Brunet *et al.*, 2004; Mejía-Roa *et al.*, 2008; Fertig *et al.*, 2012; Ochs *et al.*, 2009; Fertig *et al.*, 2013; Kossenkov and Ochs, 2009; Wang *et al.*, 2006). However, the continuous and interdependent nature of many NMF results can make biological inference challenging especially when searching for biomarkers or genetic drivers. A method to obtaining genes that uniquely identify NMF solutions would eliminate these challenges. Here, we develop PatternMarkers, a statistic to take the relative gene weights output from NMF algorithms and to return only those genes that are strongly associated with a particular pattern or with a linear combination of patterns. Identifying unbiased biomarkers using PatternMarkers requires genome-wide transcriptional data. To maximize the potential for novel marker detection, we set out to expand the O(1,000) gene limit, which is typical to achieve convergence in NMF, to the O(10,000) genes comprising the entire human transcriptome. Currently, NMF methods are highly dependent upon the genes selected or compaction methods to limit the size of the data matrices used for analysis (de Campos et al., 2013). Therefore, we developed GWCoGAPS, a whole genome implementation of CoGAPS (Fertig et al, 2010), a Markov chain Monte Carlo (MCMC) NMF that encodes sparsity in the decomposed matrices with an atomic prior (Sibisi and Skilling, 1997). Previously, we demonstrated that CoGAPS analysis of datasets containing representative subsets of the genes converge with similar patterns. These patterns can then be fixed to a consensus pattern across the datasets to provide a robust whole-genome NMF, without the prohibitively large computational cost of NMF factorization of a single matrix containing the entire genome. GWCoGAPS takes advantage of parallel computing to massively cut runtime and ensure genome-wide convergence. We also include a Shiny web application, patternMatcher, to compare patterns across parallel runs to increase robustness and interpretability of the resulting patterns. Using patternMarkers with GWCoGAPS to analyze tissues from the Genotype-Tissue Expression Project (Consortium et al., 2015), we parsed patterns of expression specific to brain regions and cell types to demonstrate the power of these algorithms for biomarker discovery.

## 2 Methods

NMF decomposes a data matrix of D with N genes as rows and M samples as columns, into two matrices, the pattern matrix P with rows associated with BPs in samples and the amplitude matrix A with columns indicating the relative association of a given gene in each BP. CoGAPS is a Bayesian NMF that incorporates both non-negativity and sparsity in A and P as described in (Fertig et al., 2010). The number of BPs (columns of A and rows of P, K) is an argument. Both the PatternMarkers statistic and GWCoGAPS are in the CoGAPS Bioconductor package as of version 3.5 and are generalized for other NMF algorithms.

The patternMarkers statistic finds the genes most uniquely associated with a given pattern or linear combination of patterns by computing

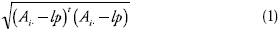
where *A*_*i*_ are the elements of the A matrix for the *i*^th^ gene scaled to have a maximum of one and *l* is the *p*^th^ user specified norm. PatternMarkers defaults to *p*=*k*, such that *l* is the identity vector and the associated distance is computed separately for each of the *k* patterns. Unique sets are generated by ranking a genes associated distances from each norm such that the higher the rank of the gene, the less it is associated with the considered pattern. Genes are subset by their lowest ranking pattern or thresholded by the first gene to have a lower ranking in another patterns.

The GWCoGAPS function automates and parallelizes the whole-genome CoGAPS analysis from Fertig et al. (2013) in a single R function. GWCoGAPS has three parameters: the number of sets for partitioning the whole genome data, the seed for each Markov Chain, and the method for determining the consensus patterns. A new modification to CoGAPS, setting the seed both ensures that each set of genes is run with a different set of random numbers and that runs on any dataset are reproducible. A default pattern matching function is provided along with a Shiny-based web application patternMatcher for recompiling the parallelized results (Supplemental Fig. 1A). Additional runtime options, input, and manual implementations are described in the GWCoGAPS vignette.

RPKM data for the seven samples with most brain regions was downloaded from dbGaP. GWCoGAPS was run for a range of k patterns with k=10 selected and uncertainty as 10% of the data (Fertig et al. 2013). The code to reproduce this analyses and the GWCoGAPS results are in Supplemental Files 3 and 4.

## 3 Results

We apply GWCoGAPS to analyze patterns related to brain regions for different individuals in GTeX. The GWCoGAPS solutions for the initial parallel runs of on of the patterns is used to illustrate the strong association between patterns identified from the subsets using patternMatcher (Supplemental Fig. 1A). The first pattern highlights GWCoGAPS ability to deconvolute tissue specific signatures (Supplemental Fig. 1B). This pattern uniquely identifies the cerebellum, determined to be the most distinct region by the consortium (Consortium et al., 2015). GTEx found that strong individual specific effects increases with tissue relatedness as illustrated by their inability to achieve tissue specific clusters of the different brain regions by expression alone (Melé et al., 2015; Consortium et al., 2015). By allowing for gene reuse across different patterns, GWCoGAPS is able to overcome these effects to isolate the cerebellums signature as confirmed by enrichment in cerebellum development and morphogensis (GO:0021549 p=2.1E-04, GO:0021587 p=3.4E-03).

The second pattern illustrates PatternMarkers power as inference is difficult from the GWCoGAPS result alone (Supplemental Fig. 1B). This pattern depicts subpopulations of cells residing in multiple brain regions derived from common precursors in the dorsal pallium. Progeny of the pallium are specified by the transcription factors TBr1 and Emx1 (Remedios et al., 2007) ranked second and fourth by the patternMarker statistic for this pattern. Gene set enrichment tests further confirms enrichment for pallium development (GO:0021543 p=1.6E-08.

Deconvolution of cell type and tissue specific signatures from aggregate data represent a major technical challenge. We have illustrated the unique ability of GWCoGAPS, the first whole genome Bayesian NMF, to accomplish this. The manual pipeline and shiny app, patternMatcher, also expanded this methodology to a variety of NMF techniques. Finally, the PatternMarkers statistic derives gene sets uniquely representative of BPs from the continuous weights of NMF solutions. Together, PatternMarkers and GWCoGAPS are a major advance in bioinformatic approaches to find data-driven biomarkers and genetic drivers in whole genome transcriptomic data.

## Funding

This work was supported by the National Institutes of Health [NCI R01CA177669 and K25CA141053 to E.J.F., NLM R01LM001100 to M.F.O. and NCI P30 CA006973] and the Cleveland Foundation and Johns Hopkins University Discovery Awards to E.J.F.

## Conflict of Interest

none declared.

---

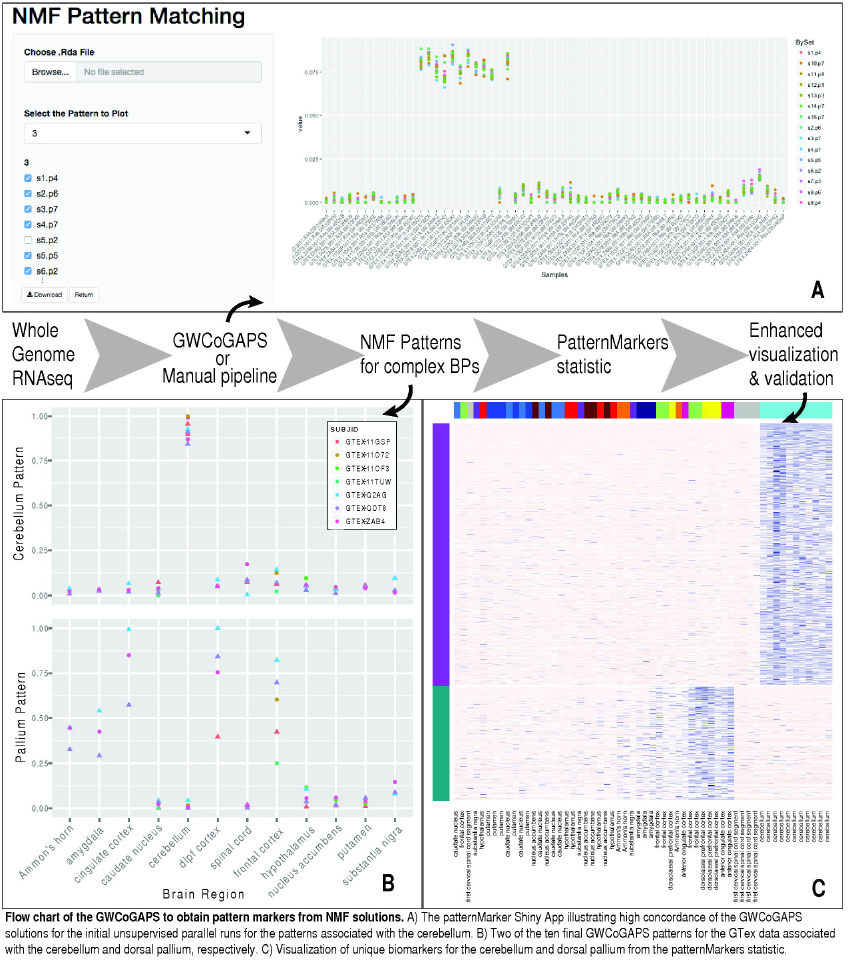

